# Quantification of microbial robustness

**DOI:** 10.1101/2021.12.09.471918

**Authors:** Cecilia Trivellin, Lisbeth Olsson, Peter Rugbjerg

## Abstract

Stable cell performance in a fluctuating environment is essential for sustainable bioproduction and synthetic cell functionality; however, microbial robustness is rarely quantified. Here, we describe a high-throughput strategy for quantifying robustness of multiple cellular functions and strains in a perturbation space. We evaluated quantifications theory on experimental data and concluded that the mean-normalized Fano factor allowed accurate, reliable, and standardized quantification. Our methodology applied to perturbations related to lignocellulosic bioethanol production showed that *Saccharomyces cerevisiae* Ethanol Red exhibited both higher and more robust growth rates than CEN.PK and PE-2, while a more robust product yield traded off for lower mean levels. The methodology validated that robustness is function-specific and characterized by positive and negative function-specific trade-offs. Systematic quantification of robustness to end-use perturbations will be important to analyze and construct robust strains with more predictable functions.

**Graphical Abstract:** 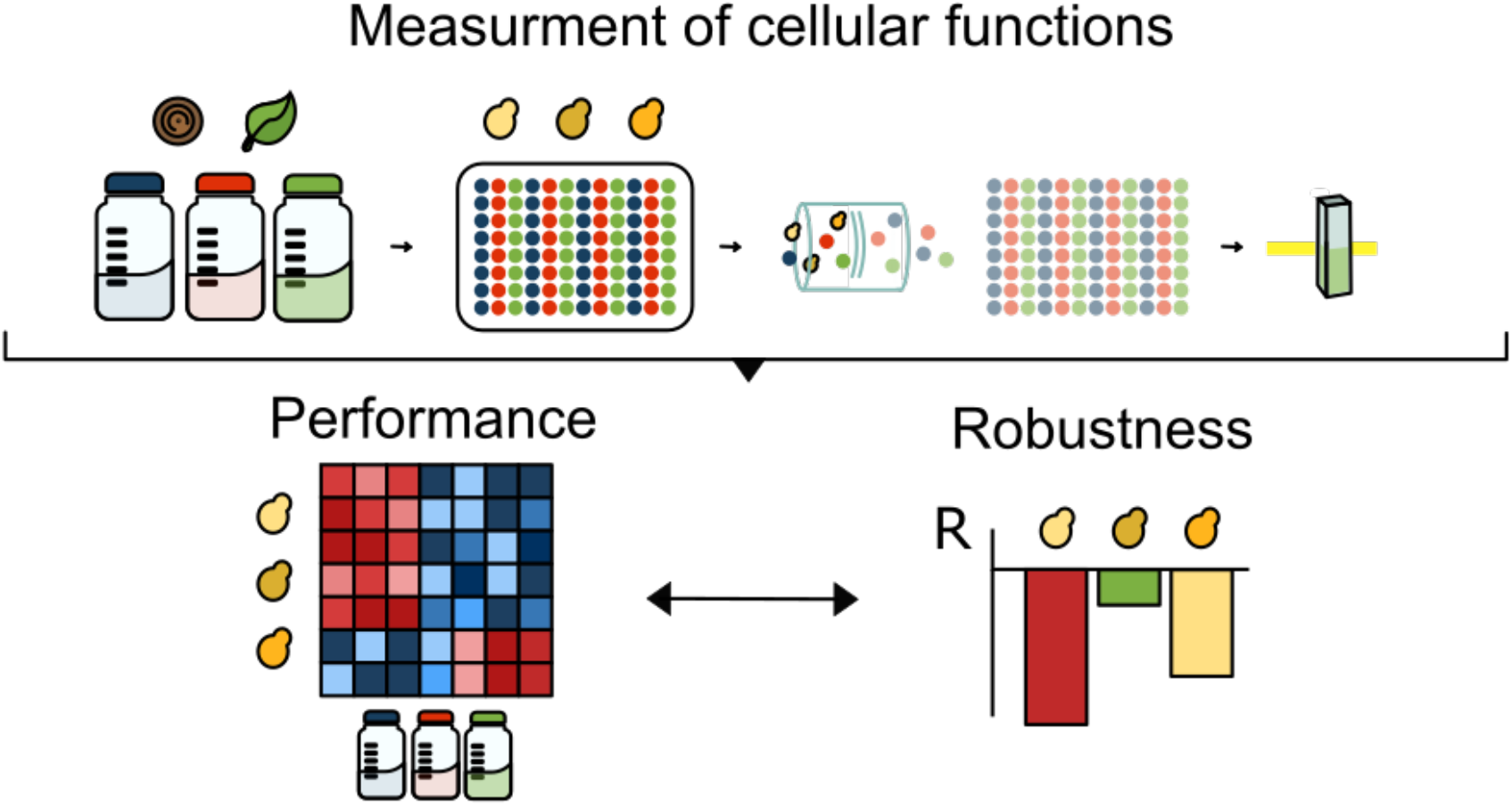

## Introduction

Microbial robustness ensures predictable, stable synthetic cellular functionality in spite of internal or external perturbations^1–3^. Robust cell manufacturing and destination performance will be a key in realizing new synthetic biology modalities and efficient bioproduction^4,5^. Robustness is defined for a specific function (or phenotype) and set of perturbations^6^. Robustness is therefore different from tolerance, which specifically describes stable growth or survival to various perturbations e.g. via specific growth rates^7^. *In silico* systems biology quantifies robustness by the influence on a cellular function of a frequency-normalized perturbation space relative to a control condition^8^ with a normally distributed mean and standard deviation. Experimentally, measures of variation represent the stability (dispersion) of quantitative traits across perturbations, but not at different scales^6^, for which the dimensionless coefficient of variation (CV) is better suited^9^. Yet even if central to realizing predictable, scalable synthetic biology, robustness is seldomly quantified experimentally for strain functions, which may be subject to genetic or environmental perturbations of stochastic or determined behavior^5,10,11^. Here, we present and validate a high-throughput methodology to experimentally and systematically quantify microbial robustness with script available from Github. We show that the methodology can be used to systematically quantify and compare the robustness of different strain functions of interest in a relevant perturbation space and relate them to their performance level. Precise quantification will allow for exploration of tradeoffs between robustness and performance of different cellular functions.

## Results and Discussion

### Development of a systematic method for quantification of robustness based on the Fano factor

To experimentally quantify robustness (R), we first set four important criteria to ensure consistency, reproducibility, and standardization. 1) Testing more perturbations should not change R, only its statistical significance. 2) Positive and negative deviations from the mean or a control performance level^8^ should contribute negatively to R. 3) Higher R should represent greater robustness. 4) R should be dimensionless and capture cellular function values at different orders of magnitude allowing comparison. To meet these criteria, we evaluated the reported theory ^8,9^. The first theory quantifies R as the CV based on standard deviation/mean 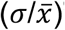^9^. R_CV_ was calculated as:

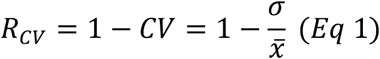

However, when cellular functions are subjected to different experimental perturbations, CV becomes > 1 complicating interpretation. More importantly, as others have, we found CV was poorly accurate in describing data dispersion with means between 0 and 1 (Figure S1)^12^. CV therefore failed our fourth criterion.

The second theory quantifies R by evaluating a function change in relation to a specific control condition 0, according to Kitano’s formula^8^:

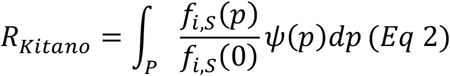

R_Kitano_ reports the ratio between the perturbed function *f_i,s_*(*p*) and the control condition *f_i,s_*(0) over a space of perturbations P, each multiplied for its frequency *ψ*(*p*). We simplified Eq2 and assumed an equal frequency for each perturbation. However, functions performing better than the control achieved higher robustness (Figure S2), further, defining a control condition performance is not always meaningful. As a result, R_Kitano_ failed our criteria 1,2 and 4.

Therefore, we evaluated an approach quantifying R as the dispersion of data around the function means using the Fano factor (Figure 1). The Fano factor is commonly used to study transcriptional bursting and noise in gene expression by identifying deviation from Poissonian behavior and has been proposed for robustness before ^6,12–14^, but not actually deployed. For each function *i*, a strain *S*, and a perturbation space *P*, R_S,i,P_ was calculated as 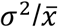 ^15^:

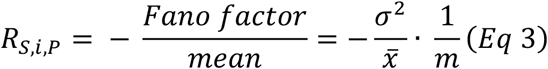

**Figure 1:**
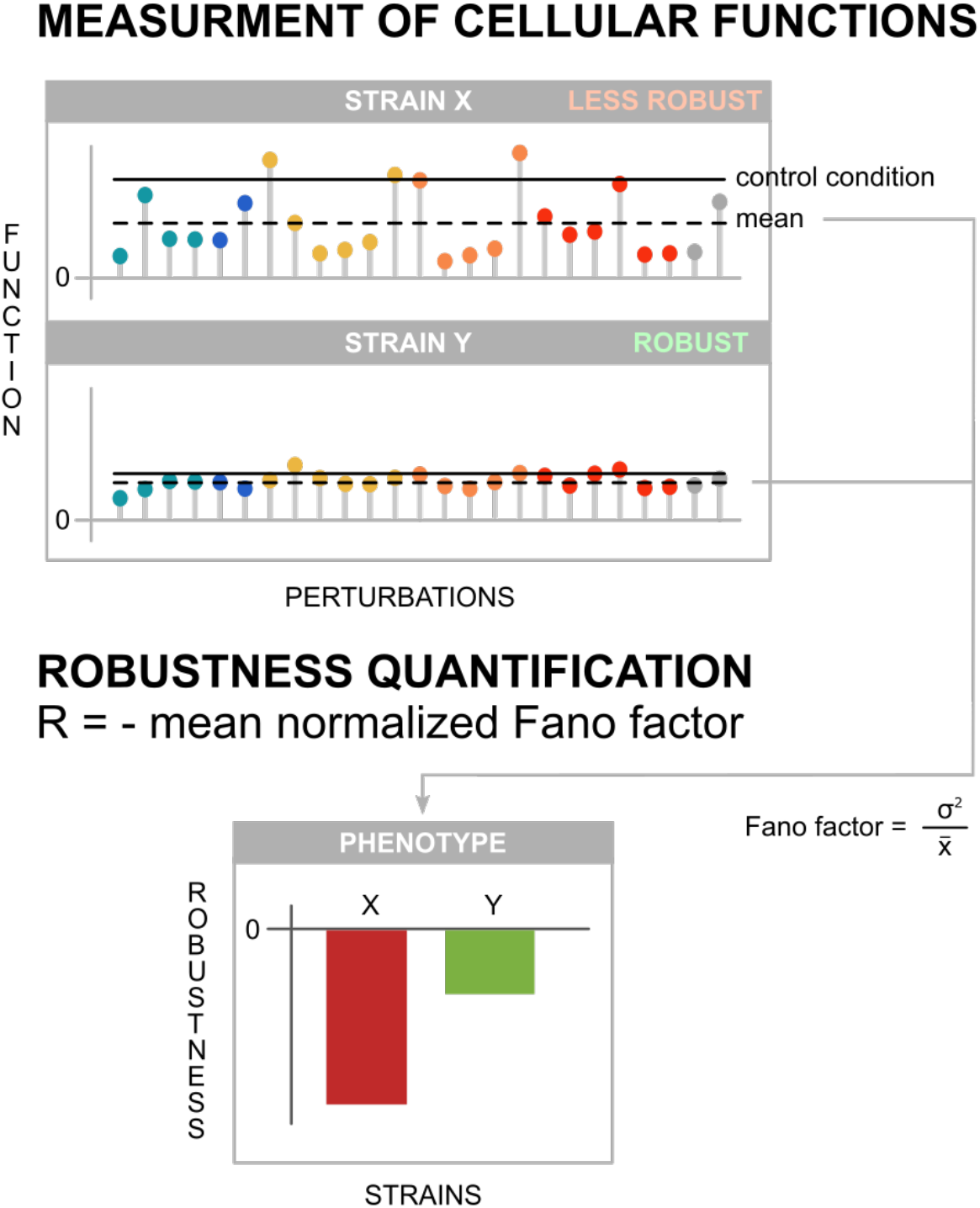
Relevant functions are measured upon exposure to various perturbations (colored dots) and robustness is calculated as the negative mean-normalized Fano factor. A control condition (e.g. 20 g/L glucose) is needed for calculation of R_Kitano_.

To allow comparison between functions, we normalized the Fano factor to the mean *m* of all strains (Figure 1). We set the upper limit for R to 0 (highest robustness) and the problem of 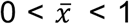 was solved. As the Fano factor remained finite for means approaching zero, the weight of the mean on R was higher than for the CV. This quantification strategy was frequency independent, dimensionless, and free from arbitrary control conditions, thereby meeting all criteria.

### Quantifying robustness of five cellular functions: Case study of lignocellulosic bioethanol production

To validate the methodology for quantifying microbial robustness, we used lignocellulosic bioethanol production (see Supporting Information, Material and methods). We included the *Saccharomyces cerevisiae* CEN.PK113-7D laboratory strain^16^, and the industrial strains Ethanol Red and PE-2, whose robust ethanol production and growth is advantageous in starch and sugarcane fermentation^17–19^. We measured five relevant cellular functions (maximum specific growth rate, lag phase, cell dry weight, biomass, and ethanol yields) across 29 different perturbations (single-component lignocellulose growth conditions) in a 96-well plate high-throughput setup (Methods and materials).

The three strains exhibited different production and growth functionality when exposed to the 29 perturbations (Figure 2A). Across the lignocellulose perturbation space, Ethanol Red performed better than CEN.PK and PE-2 in all functions, except for ethanol yield. Aldehydes had a negative effect on all five measured functions, while pentoses resulted in unchanged or improved functions. Lactic, levulinic, formic and acetic acid reduced cell dry weight and biomass yield (Figure 2A).

**Figure 2:**
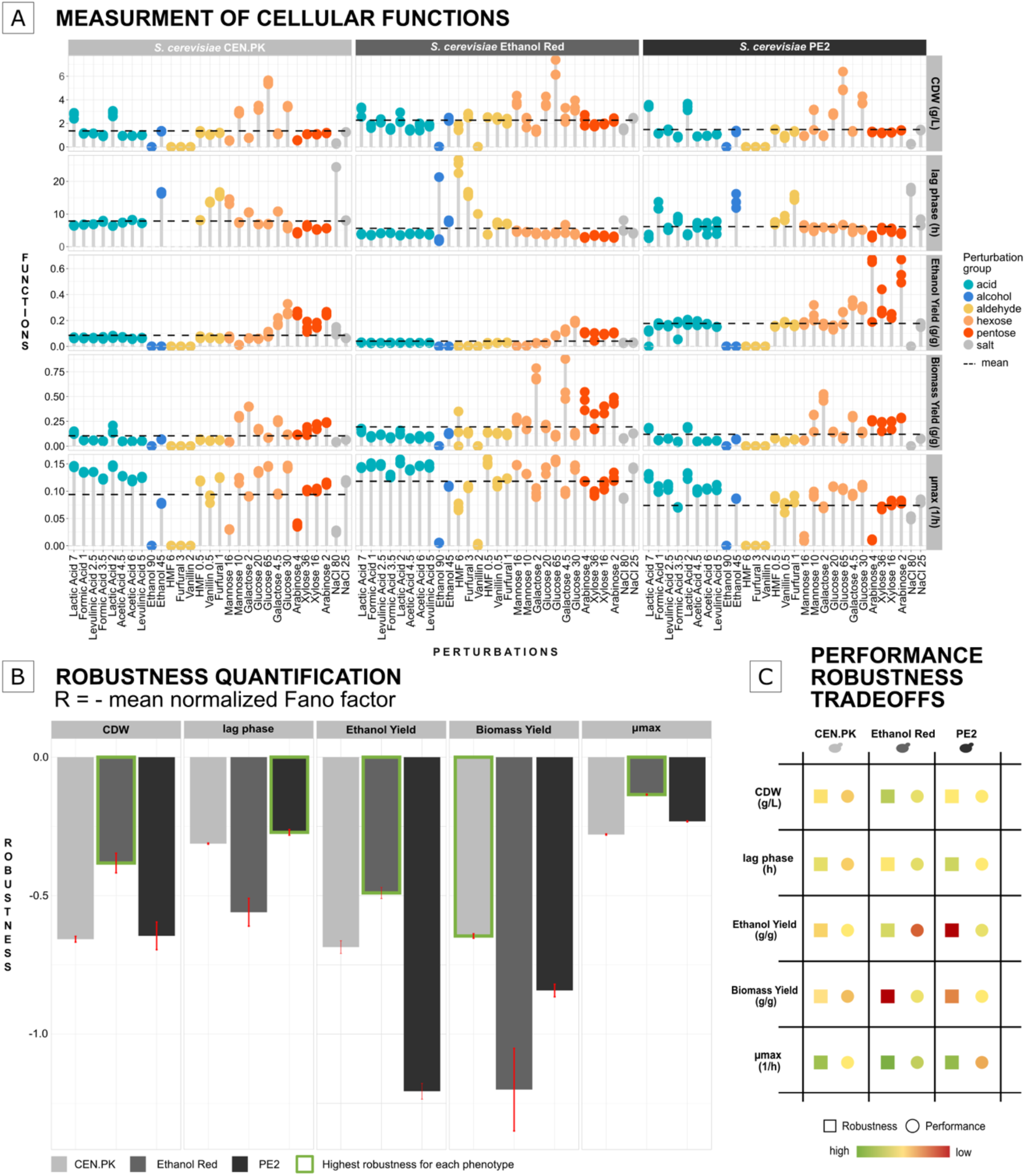
Quantification of microbial robustness with the Fano factor. (A) Function evaluation in a large perturbation space containing components found in lignocellulosic hydrolysates. CDW: cell dry weight; μmax: maximum specific growth rate. All points are individual biological replicates (n = 3). Lag phase missing points: cultures did not grow within 48 hr. (B) Robustness quantification for each function. Error bars: standard error of the mean (n = 3). (C) Robustness and performance trade-offs for each function and strain.

We next quantified the robustness and found the maximum specific growth rate, ethanol yield, and cell dry weight as significantly higher in Ethanol Red (cell dry weight (p-value < 0.005), maximum specific growth rate (p-value < 7e-8), ethanol yield (p-value <0.001)) (t-test) (Figure 2B), supported by data less dispersed around the mean. These robust cellular functions were accompanied by a more fragile biomass yield, most robust in CEN.PK (p-value <0.02), and lag phase most robust in PE-2 (p-value < 0.002) (Figure 2B). PE-2 achieved high mean ethanol yield, but its robustness was the lowest in part due to positive effects from pentoses. The observed very robust growth and production functions of Ethanol Red could explain its application in first-generation bioethanol plants that share our perturbations mainly except from aldehydes^18,^, but came with a cost of lower average performance. High performance and high robustness are sometimes considered mutually exclusive^20^. In theory, these two properties are believed to trade off with one another in several biological systems^21,22^. Our quantification methodology identified possible robustness and performance trade-offs in lignocellulose-based bioethanol production (Figure 2C). For example, Ethanol Red traded its ethanol yield performance for high robustness, and vice versa for PE-2. Oppositely, we found that PE-2 traded robustness for performance for its maximum specific growth rate. Based on these findings, we observed that correlations between performance and robustness are both function and strain depended. Larger datasets are needed to establish such correlations.

A systematic framework for assessing robustness of several performance indicators will improve our understanding of how cellular functions respond to relevant perturbations, e.g., by favoring robustness, performance, or a sub-optimal state for both. In strain engineering, robust yields of products are sometimes preferred over higher but unstable yields^5^. In synthetic biology, engineered strains should carry new functions that perform robustly under anticipated perturbations. Relevant functions include biosensor signals, gene expression reporters and heterologous proteins production. Quantifying robustness to scale-up or long-term cultivation (e.g. genetic robustness) will also be highly relevant to prevent declines of performance e.g. due to accumulation of mutations and heterogeneity^5,23^ e.g. by quantifying robustness over many cellular divisions.

High-throughput-friendly methodology as described here will be useful for quickly quantifying robustness of multiple strain variants and functions. The methodology can uncover which perturbations affect the robustness of specific functions, how robustness relates positively and negatively to performance for each function (i.e. trade-offs). Through our methodology we validated that robustness is function-specific^6^, rather than universal strain value, as previously theorized.

Future work will also show whether more robust starting strains are better for subsequent enhancements via metabolic or evolutionary engineering. Robustness quantification may also include perturbation frequency, as well as more strains and perturbations. At present, our methodology may be applied to phenomics databases to compare numerous traits and strains.

## Supporting Information

Figure S1: Robustness estimation with Eq1.

Figure S2: Robustness estimation with Eq2.

Material and methods.

## Author information

### Notes

The developed scripts for robustness quantification are available at Github: https://github.com/cectri/Quantification-of-microbial-robustness

## Acknowledgments

Financial support by Novo Nordisk Foundation grant DISTINGUISHED INVESTIGATOR 2019 - Research within biotechnology-based synthesis & production **(#0055044**) is gratefully acknowledged.

Société Industrielle Lesaffre, Division Leaf, is kindly acknowledged for providing the Ethanol Red strain.

