## Supporting information for "Quantification of microbial robustness"

### Figure S1

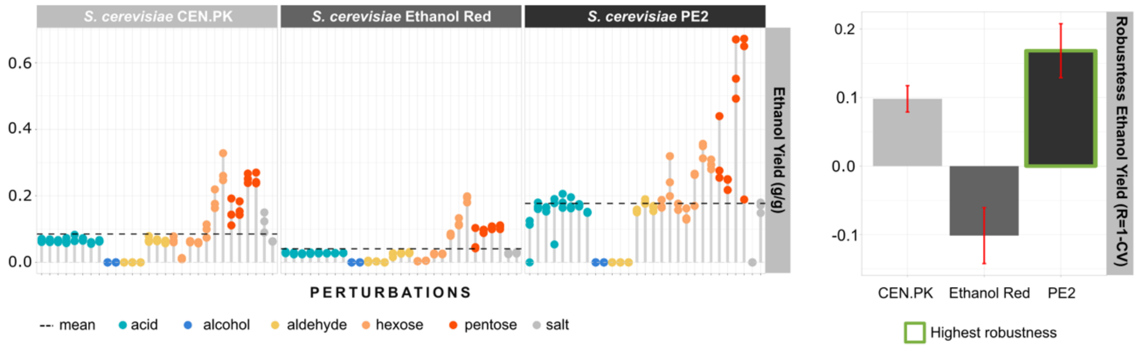


*Figure S1: Robustness estimation with Eq1. The three panels to the left denote different S. cerevisiae strains grown in medium containing various components (grouped by color) that mimic lignocelluloses hydrolysates. Ethanol yield (g/g) measurements from the database created in the case study (see Material and methods) are reported. The plot to the right shows robustness of the ethanol yield presented in the left panels. Robustness (R) was calculated with Eq1 based on the coefficient of variation (CV) by applying the formula R = 1 - CV. Error bars correspond to the standard error of the mean.*

### Figure S2

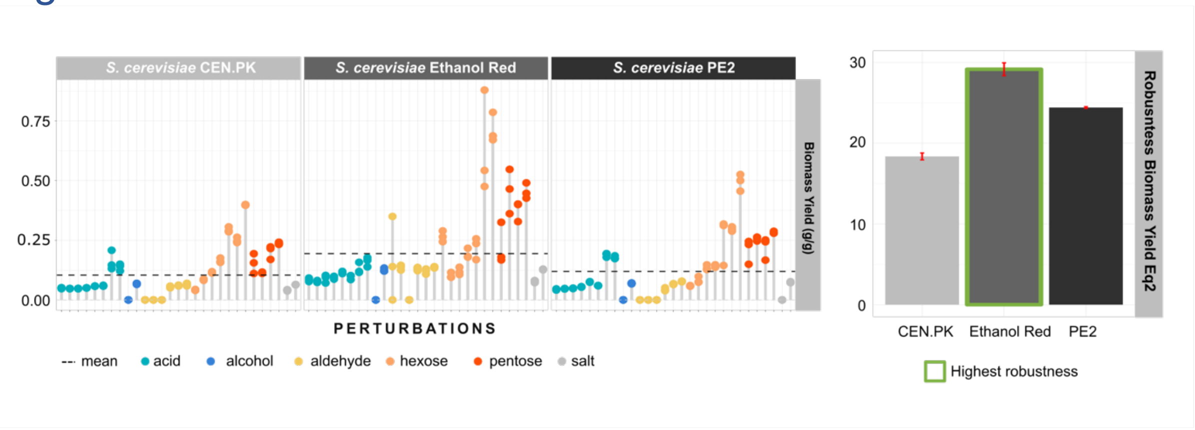


*Figure S2: Robustness estimated with Eq2. The three panels to the left denote different S. cerevisiae strains grown in medium containing various components (grouped by color) that mimic lignocelluloses hydrolysates. Biomass yield (g/g) measurements from the database created in the case study (see Material and methods) are reported. The plot to the right shows robustness of the biomass yield presented in the left panels. Robustness was calculated with Eq2. Error bars correspond to the standard error of the mean.*

### Material and methods

#### *Strains*

The strains used in the study were *Saccharomyces cerevisiae* CENPK113-7D[1] (Scientific Research and Development GmbH, Oberursel, Germany), PE-2[2] (wild-type strain isolated during sugarcane-to-ethanol production in Brazil), and Ethanol Red® (kindly provided by Société Industrielle Lesaffre, Division Leaf).

#### *Media preparation*

Delft minimal medium[3] was used for strain cultivation. The medium was prepared with 5 g/L $(NH_4)_2SO_4$, 3 g/L $KH_2PO_4$, 1 g/L $MgSO_4{\cdot}7H_2O$, 1 mL (in 1 L solution) trace mineral solution (Table S1), and 1 mL (in 1 L solution) vitamin solution (Table S2). The medium was adjusted to pH 5 with KOH and buffered with 250 mM potassium hydrogen phthalate. Multiple compounds were added to the minimal medium to mimic the composition of lignocellulosic hydrolysates obtained mostly from spruce, corn starch and wheat straw (Table S3).

*Table S1: Mineral solution composition.*

| **Chemical** | **Amount (g/L)** |
|---|---|
| EDTA | 0.015 |
| $ZnSO_4{\cdot}7H_2O$ | 0.0045 |
| $MnCl_2{\cdot}4H_2O$ | 0.0008 |
| $CoCl_2{\cdot}6H_2O$ | 0.0003 |
| $CuSO_4{\cdot}5H_2O$ | 0.0003 |
| $Na_2MoO_4{\cdot}2H_2O$ | 0.0004 |

| | |
|---|---|
| $CaCl_2 \cdot 2H_2O$ | 0.0045 |
| $FeSO_4 \cdot 7H_2O$ | 0.003 |
| $H_3BO_3$ | 0.001 |
| KI | 0.0001 |

*Table S2: Vitamin solution composition.*

| **Vitamin** | **Amount (g/L)** |
|---|---|
| d-Biotin | 0.00005 |
| Calcium D(+) pantothenate | 0.001 |
| Nicotinic acid | 0.001 |
| Myo-inositol | 0.025 |
| Thiamine HCl | 0.001 |
| Pyridoxine HCl | 0.001 |
| Para-aminobenzoic acid | 0.0002 |

*Table S3: List of chemicals (and their relative concentrations) added to the medium to mimic the composition of lignocellulosic hydrolysates.*

| **Chemical** | **Carbon source** | **Concentration (g/L)** | **Reference** |
|---|---|---|---|
| D-(+)-Glucose | | 65, 30, 20 | [4,5] |
| D-(+)-Xylose | D-(+)-Glucose (5 g/L) | 36, 16 | [4] |
| D-(+)-Galactose | | 4.5, 2 | [5,6] |
| L-(+)-Arabinose | D-(+)-Glucose (5 g/L) | 4, 2 | [6,7] |
| D-(+)-Mannose | | 16, 10 | [5,6] |
| Formic acid | D-(+)-Glucose (20 g/L) | 3.5, 1 | [4–6] |

| | | | |
|---|---|---|---|
| Acetic acid | D-(+)-Glucose (20 g/L) | 6, 4.5 | [4,6,7] |
| DL-Lactic acid | D-(+)-Glucose (20 g/L) | 7, 2 | [8,9] |
| Levulinic acid | D-(+)-Glucose (20 g/L) | 5, 2.5 | [5,6] |
| 5-Hydroxymethylfurfural | D-(+)-Glucose (20 g/L) | 6, 0.5 | [4,5] |
| Furfural | D-(+)-Glucose (20 g/L) | 3, 1 | [4,6] |
| Vanillin | D-(+)-Glucose (20 g/L) | 2, 0.5 | [6] |
| Ethanol | D-(+)-Glucose (20 g/L) | 90, 45 | [10] |
| NaCl | D-(+)-Glucose (20 g/L) | 80, 25 | [11] |

##### *Fermentation experiments*

The strains were preserved in glycerol (final concentration 16%) at -80°C. Pre-cultures were prepared by inoculating 10 μL glycerol stocks in 5 mL of the above-described Delft medium (20 g/L glucose), followed by incubation at 30°C with 200 rpm shaking.

The optical density at 600 nm ($OD_{600}$) of overnight cultures was monitored in a GENESYS™ 10 spectrophotometer (Thermo Scientific). After 24 h, the cultures (in exponential phase) were inoculated in square polystyrene 96-half-deepwell microtiter plates (CR1496dg; Enzyscreen) at a starting $OD_{600}$ of 0.02. Strain growth was monitored in a Growth Profiler 960 (Enzyscreen) and was expressed as green value (GV) units. The microplates were covered with a $CO_2$-release cover (CR1296t; Enzyscreen) to minimize passive diffusion of $O_2$ and mimic anaerobic conditions. Each plate was used for a single strain, each well corresponded to a specific growth condition, and each condition was assayed in three technical replicates. The strains were cultivated in the growth profiler for 48 h at 30°C, with 250 rpm shaking. After 48 h, the plates were removed from the growth profiler and the GV units at 48 h were converted to OD values according to the following formula:

$$OD = a(GVvalue - GVblank)^b + c(GVvalue - GVblank)^d + e(GVvalue - GVblank)^f$$

The constants were as follows: $a$ = 0.019, $b$ = 1; $c = 3.82 \times 10^{-6}$, $d$ = 2.66, $e = 3.111 \times 10^{-22}$, $f$ = 10.5, and $GVblank$ = 26.3.

The equation was previously calibrated using Delft medium and *S. cerevisiae* CEN.PK113-7D. Cultures were diluted based on the final $OD_{600}$ measured in the growth profiler. $OD_{600}$ of the culture at 48 h was measured in a plate reader (SPECTROstar nano; BMG LABTECH). The culture was transferred on a hydrophilic polytetrafluoroethylene multiscreen solvinert 96-well filter plate (MSRLN0410; Millipore) and filtered into a new 96-well microtiter plate (82.1581; Sarstedt).

##### *Cell dry weight (CDW) determination*

*S. cerevisiae* strains were cultivated in Delft medium (20 g/L glucose) to stationary phase and centrifuged at 5000 rpm for 5 min. The cell pellet was resuspended in 1 mL water and $OD_{600}$ was measured. A series of five 1:2 dilutions were made and filtered through a pre-dried and weighed 0.45-µm polyethersulfone membrane (Sartorius). $OD_{600}$ of the dilutions was measured. The filters containing the samples were dried for 10 min in a microwave oven (350 W) and weighted again. Calibration curves were constructed with CDW and $OD_{600}$. The slope values for each strain were used subsequently to calculate the CDW of the growth profiler cultures.

##### *Determination of sugars and ethanol*

Culture medium obtained by filtration from the first cultivation was used to determine the sugars and ethanol content. Ethanol was measured with the K-ETOH Ethanol Assay Kit, glucose with the K-GLUHK-220A D-Glucose HK Assay Kit, mannose with the K-MANGL D-Mannose/D-Fructose/D-Glucose Assay kit, xylose with the K-XYLOSE D-Xylose Assay Kit, and galactose and arabinose with the K-ARGA L-Arabinose/D-Galactose Assay Kit (all Megazyme). The assays are based on enzymatic reactions, which produce NADH, whose absorbance at 340 nm can be read in a spectrophotometer. The amount of NADH is stoichiometric with the amount of the compound of interest, making it possible to calculate the concentration (g/L) of the respective compounds.

##### *Determination of performance values.*

The growth data in GV units were imported in R software for visualization and determination of growth parameters. The maximum specific growth rate was determined using all_*splines* function. Duration of the lag phase was determined by calculating the x coordinate of the intersection between the line with µmax slope passing through the inflection point and the line passing through y0 parallel to the x-axis. In the wells where no growth was detected, $R^2$ was < 0.99, so the µmax was set to 0 while the lag phase was set to NA.

Ethanol yield and biomass yield were calculated based on total consumed sugars as follows:

$$Ye = \frac{ethanol\ produced\ (g)}{initial\ sugars\ (g) - final\ sugars\ (g)} \text{ (Eq1)}$$

$$Yb = \frac{CDW\ (g)}{initial\ sugars\ (g) - final\ sugars\ (g)}\ \text{(Eq2)}$$

Scripts with line-by-line explanation available on Github (https://github.com/cectri/Quantification-of-microbial-robustness).

##### *Robustness calculation*

Eq3 was applied to the case study database mentioned above. The following variables were considered: μmax (1/h), lag phase (h), CDW (g/L), biomass yield (g biomass/g consumed substrate), and ethanol yield (g produced/g consumed). Robustness and performance values were calculated and plotted in R. Statistical difference among the different cellular functions and strains was determined with an unpaired, two-sided t-test and p-values were adjusted with Holm-Bonferroni method. Scripts with line-by-line explanation available on Github (https://github.com/cectri/Quantification-of-microbial-robustness).
